# Lysosomal Glucocerebrosidase is needed for ciliary Hedgehog signaling: A convergent pathway to Parkinson’s disease

**DOI:** 10.1101/2025.01.20.633968

**Authors:** Sreeja V. Nair, Ebsy Jaimon, Ayan Adhikari, Jonas Nikoloff, Suzanne R. Pfeffer

## Abstract

Mutations in *LRRK2* and *GBA1* are the most common genetic causes of familial Parkinson’s disease. Previously, we showed that pathogenic *LRRK2* mutations inhibit primary cilia formation in rare interneurons and astrocytes of the mouse and human dorsal striatum. This blocks Hedgehog signaling and reduces synthesis of neuroprotective GDNF and NRTN, which support neurons vulnerable in PD. Here we show that *GBA1* mutations also impair Hedgehog signaling through a distinct mechanism. Loss of *GBA1* activity decreases accessible cholesterol in primary cilia of cultured cells, thereby disrupting Hedgehog signaling. In the mouse striatum, *Gba1* mutations result in reduced Hedgehog-induced *Gdnf* RNA expression in cholinergic interneurons, despite having no detectable impact on cilia formation. Also, both *Lrrk2* and *Gba1* mutations suppress Hedgehog-induced *Bdnf* expression in striatal astrocytes. These findings underscore the role of Hedgehog signaling in the nigrostriatal circuit and reveal a convergent mechanism by which distinct mutations may contribute to PD pathogenesis.

## INTRODUCTION

Pathogenic mutations in the β-glucocerebrosidase (*GBA1*) gene cause Gaucher disease, a recessive lysosomal storage disorder that results in accumulation of glucocerebroside and glucosylsphingosine in the liver, spleen and bone marrow^1^. For reasons that are not understood, homozygous and heterozygous carriers of *GBA1* mutations are at twenty-fold increased risk for developing Parkinson’s disease (PD)^1–4^, and mutations in *GBA1* are now recognized to be the greatest known genetic risk factor for development of PD^5^. *GBA1*-PD is similar to idiopathic PD but with earlier onset of motor symptoms, and higher rates and earlier appearance of cognitive decline.

In its early stages, PD is associated with specific loss of dopamine neurons in the substantia nigra. Yet, loss of GBA1 activity would be predicted to cause general lysosome dysfunction and would be expected to impact not only this, but also most other brain regions. Thus, additional mechanisms and pathways beyond general lysosome dysfunction are needed to explain the specific and selective link between *GBA1* mutations and dopamine neuron loss in *GBA1*-linked PD. Dominant, activating mutations in the Leucine Rich Repeat Kinase 2 (*LRRK2*) gene also cause PD, and surprisingly, PD patients with mutations in both *LRRK2* and *GBA1* genes may have a milder disease course than *GBA1*-PD patients in terms of motor function and olfaction^6,7^. These findings strongly suggest an overlap in the mechanisms by which *GBA1* and *LRRK2* mutations cause PD.^8^

Dopamine neurons of the substantia nigra project to the striatum. In the striatum, the dopamine neurons release the Sonic Hedgehog (Shh),^9^ which triggers production of neurotrophic GDNF^9–11^ and GDNF-related Neurturin (NRTN)^12^ by rare cholinergic and Parvalbumin interneurons, respectively. We have shown that pathogenic *LRRK2* mutations block formation of the primary cilia – critical compartments of Hedgehog signaling – specifically in these cholinergic and Parvalbumin interneurons (and astrocytes) in the mouse and human dorsal striatum^11–14^. Loss of cilia precludes these neurons from sensing Hedgehog signals, thus blocking their ability to produce neurotrophic GDNF^9–11^ and NRTN^12^. In the absence of neuroprotective and trophic factors, dopamine neurons that project to this brain region are left more vulnerable to cell stress and are slowly lost, consistent with a disease of aging^9,11–13,15^. In addition, in mice, we detect decreased density of fine dopaminergic neuron processes, that is likely to reflect early-stage loss of dopamine production^15^.

Cells go to great lengths to regulate cholesterol levels in specific cellular membranes^16^. While the plasma membrane contains ∼45% mol/mol cholesterol, the endoplasmic reticulum contains less than 10%. It is now appreciated that there are distinct pools of cholesterol at the plasma membrane: a chemically available, “accessible” pool needed for ciliary Hedgehog signaling, a sphingolipid-sequestered pool, and a third pool that is sequestered by membrane proteins and is essential for membrane integrity^17^. Cells obtain cholesterol by biosynthetic processes or by low density lipoprotein (LDL) endocytosis^18^. LDL-derived cholesterol is transported from inside lysosomes to the cell surface by a process that begins with the lumenal NPC2 protein moving endocytosed cholesterol from intralumenal vesicles to the NPC1 transporter for export from lysosomes^19^. In cells in which NPC proteins are absent or non-functional, cholesterol accumulates in lysosomes and cells respond to apparent cholesterol starvation by turning on transcriptional pathways of cholesterol biosynthesis^20,21^. Cholesterol accumulation in lysosomes is accompanied by accumulation of glycosphingolipids^22,23^, which may be attributed to cholesterol sequestration of these lipids upon their intra-lysosomal accumulation in NPC-deficient cells.

We recently reported that NPC1-deficient cells show increased accessible cholesterol in lysosomes, accompanied by decreased, total cell surface-accessible cholesterol levels^24^. We show here that chemical inhibition NPC1 is accompanied by loss of ciliary accessible cholesterol and thus decreased Hedgehog signaling. Moreover, we show that genetic or chemical inhibition of GBA1 also lead to accumulation of accessible cholesterol in lysosomes, decreased ciliary accessible cholesterol, and defects in Hedgehog signaling in cell culture and in *Gba1* mutant mouse brain. Similar to *Lrrk2* mutant striatum, *Gba1* mutant striatum shows loss of neuroprotective factor production and decreased fine dopaminergic processes. These findings have important implications for Parkinson’s disease etiology and provide a possible convergent pathway for Parkinson’s disease caused by mutations in *GBA1* and *LRRK2* genes.

## RESULTS

### NPC1 inhibition decreases ciliary accessible cholesterol and Hedgehog signaling

We showed previously that accumulation of cholesterol and glycosphingolipids in NPC1-inhibited lysosomes decreases plasma membrane accessible cholesterol levels^24^. We therefore predicted that NPC1 inhibition would be accompanied by a decrease in ciliary accessible cholesterol because the plasma membrane is continuous with the ciliary membrane. We triggered cilia formation in NIH-3T3 cells stably expressing a fluorescent primary cilia protein Somatostatin Receptor 3 (SSTR3-mApple) using serum starvation. After 24 hours of serum starvation, we were able to readily detect accessible cholesterol in primary cilia by labeling fixed cells with mNeon-ALOD4, a fluorescently labeled toxin-derived protein that binds selectively to accessible cholesterol^16,25,26^ (Figure 1A). In contrast, 72-hour inhibition of NPC1-mediated lysosomal cholesterol export with U18666A led to a dramatic decrease in surface-accessible cholesterol and a lower proportion within contiguous primary cilia (Figs. 1A,B). Ciliary accessible cholesterol is required for activation of the Smoothened protein that transduces Hedgehog signals into cells^27,28^. We also found that Hedgehog-responsive *GLI1 and PTCH1* RNA levels were significantly decreased in NPC1-inhibited NIH-3T3 cells (Figure 1C). These results indicate that cholesterol accumulation in lysosomes of NPC1-deficient cells leads to a reduction in accessible cholesterol at the cell surface and in primary cilia, thereby inhibiting Hedgehog signaling.

**Figure 1:**
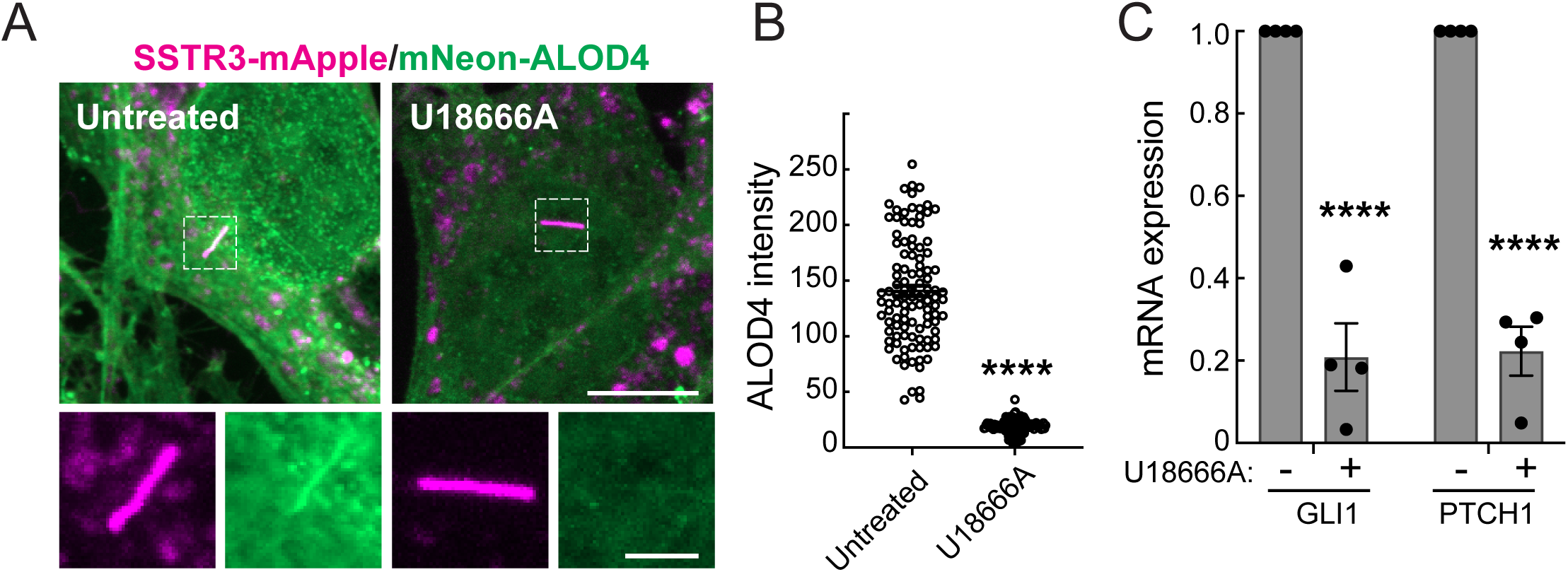
Redistribution of cholesterol in NPC1-inhibited cells limits accessible cholesterol in primary cilia and inhibits Hedgehog signaling. (A) NIH-3T3 fibroblasts stably expressing SSTR3-mApple (magenta), grown on coverslips, were treated without serum ± 1µM U18666A for 24h and fixed cells were labeled with mNeon-ALOD4 (green) to detect ciliary accessible cholesterol. Magnification bar, 10µm; inset, 2µm. (B) Quantification of ciliary mNeon-ALOD4 mean intensity from (A). Error bars represent SEM from three independent experiments with >28 cilia per condition; each dot represents one cilium. Significance was determined using Student’s t-test. ****P < 0.0001. (C) Quantification of *Gli1* and *Ptch1* RNA levels (normalized to a *Gapdh* control) from NIH-3T3 fibroblasts treated ± 1µM U18666A for 72h, with 10nM Shh in medium without serum for the last 24h. Error bars represent SEM from four independent experiments. Significance was determined relative to the control (-U18666A) by Student’s t-test. ****P < 0.0001.

### GBA1 inhibition decreases ciliary accessible cholesterol and Hedgehog signaling

In NPC1-deficient cells, both cholesterol and glycosphingolipids accumulate in lysosomes^22,23^. Since GBA1-deficient cells accumulate glucocerebroside and glucosylsphingosine in lysosomes^1^, we reasoned that these accumulated substrates might also sequester cholesterol, to effect depletion of the plasma membrane pool. To test this, we compared lysosome-accessible cholesterol in control human fibroblasts and patient-derived fibroblasts harboring various *GBA1* mutations and combinations of mutations, including N370S, the most common *GBA1* mutation among Ashkenazi Jews, and *GBA1* L444P, which is more prevalent in Asians and Caucasians of non-Ashkenazi ancestry. Consistent with a previous report,^29^ we found significant accumulation of accessible cholesterol in all *GBA1* mutant patient fibroblasts compared with controls (Figure 2A,B).

**Figure 2:**
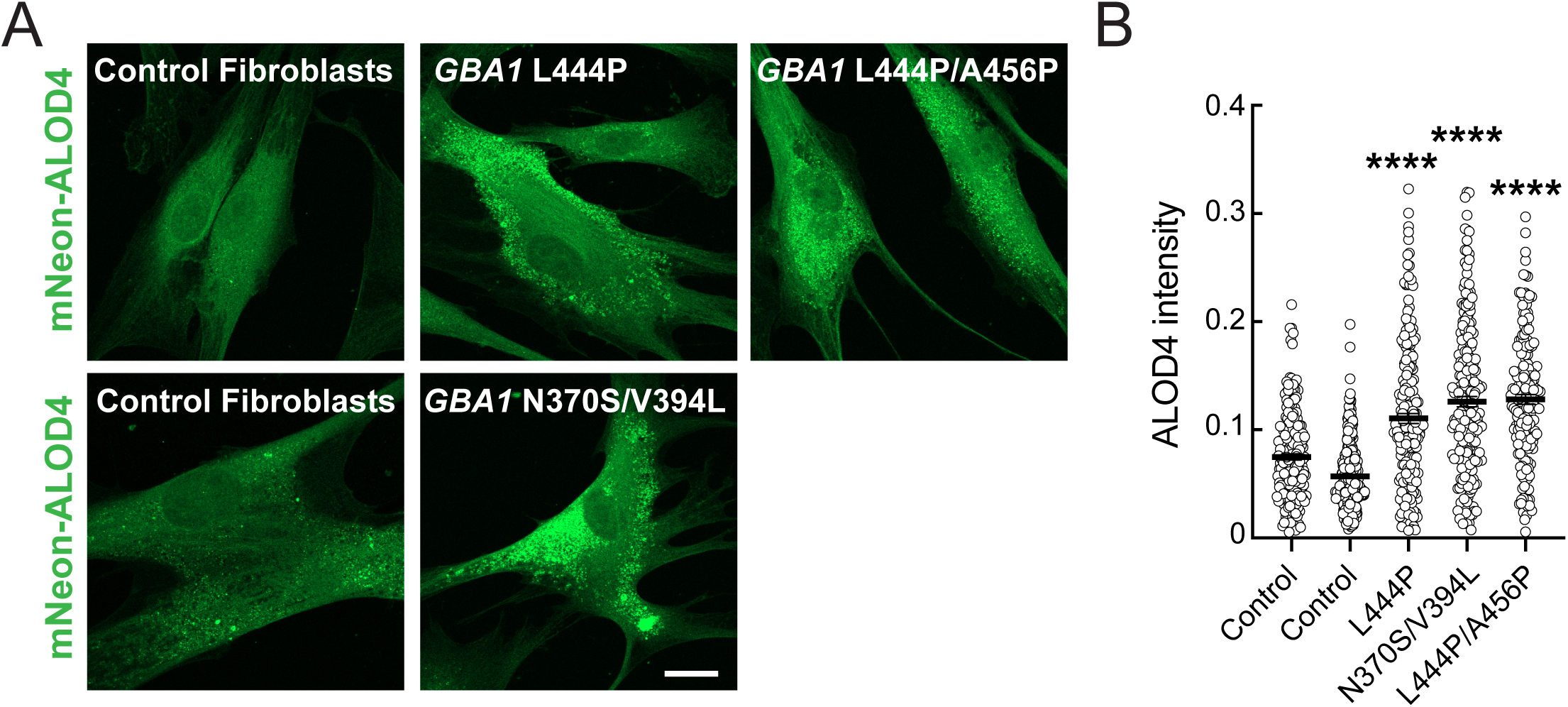
Accessible cholesterol accumulates in human *GBA1* mutant lysosomes. (A) Patient-derived *GBA1* mutant and control human fibroblasts were stained for intracellular accessible cholesterol using mNeon-ALOD4 (green). *GBA1* mutations are indicated. Magnification bar, 20µm. (B) mNeon-ALOD4 intensity per cell area from cells labeled as in (A). Error bars represent SEM from three independent experiments with >208 cells per cell line. Significance was determined relative to either set of control fibroblasts by one-way ANOVA. ****P < 0.0001.

Primary human cells bearing these *GBA1* mutations were poorly ciliated in vitro so we turned to other models to analyze the effect of GBA1-deficiency on ciliary cholesterol. NIH-3T3 cells were treated for 7 days with Conduritol beta epoxide (CBE), a small-molecule inhibitor of GBA1. GBA1 inhibition led to a significant decrease in ciliary accessible cholesterol, measured as in Figures 1 and 2 using fluorescent ALOD4 protein (Figure 3A,B). Thus, lysosomal lipid accumulation decreases accessible cholesterol at the cell surface in both NPC1- and GBA1-inhibited cells. The decrease in ciliary accessible cholesterol in cells treated with the GBA1 inhibitor was less than that seen with an NPC1 inhibitor. This is not surprising, given that cholesterol can still exit the lysosome in *GBA1* mutant cells due to continued NPC1-mediated export.

**Figure 3:**
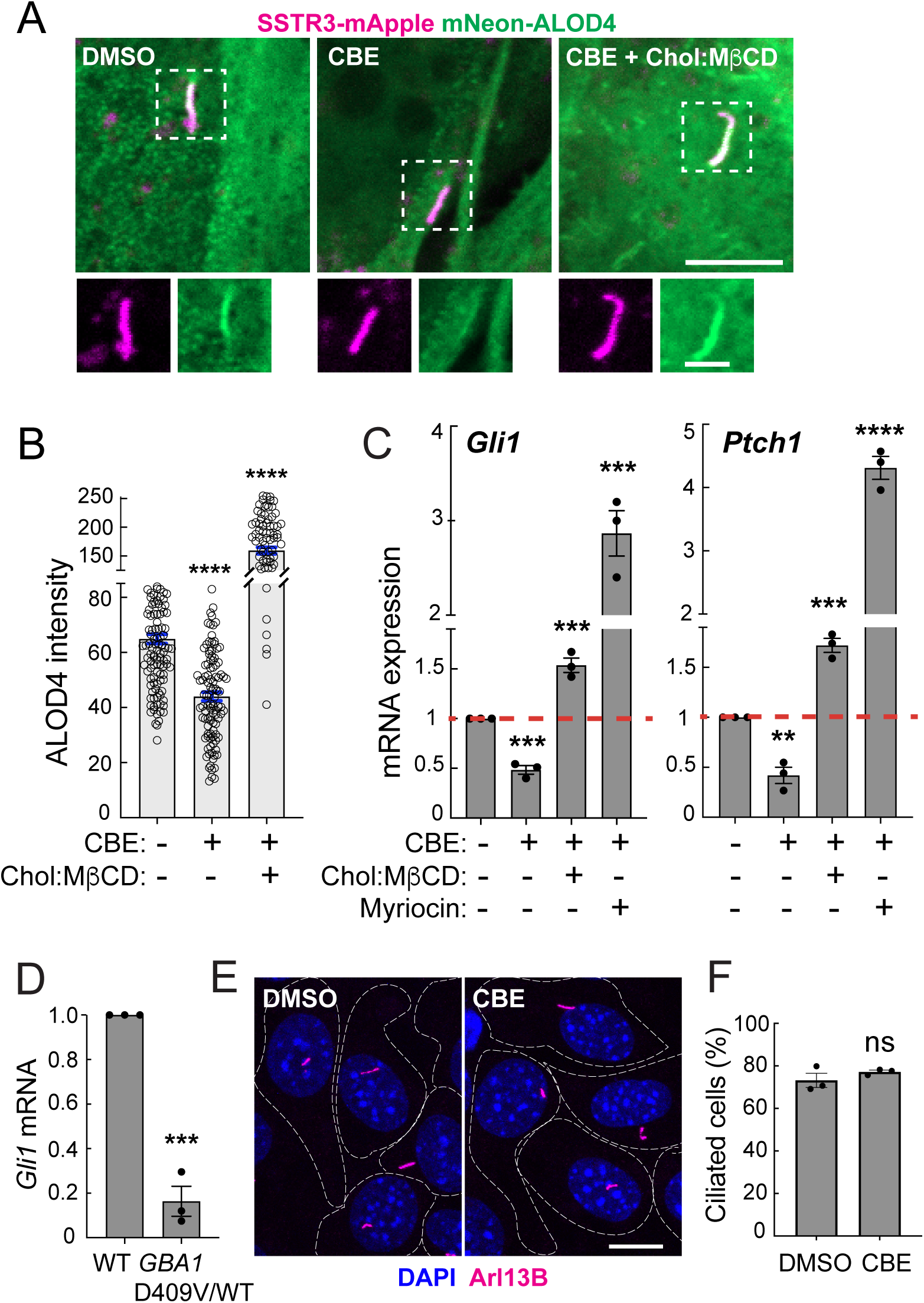
Loss of lysosomal GBA1 activity limits accessible cholesterol in primary cilia and reduces ciliary Shh signaling. (A) NIH-3T3 fibroblasts were treated with vehicle (DMSO) or 100 µM CBE for 7d. On the sixth day, media was replaced with serum-free media ± CBE, ± 0.3 mM Cholesterol:MβCD complex. Fixed cells were stained for ciliary accessible cholesterol using mNeon-ALOD4. Magnification bar, 5µm; insets (boxed regions), 2µm. (B) Quantification of ciliary accessible cholesterol from images as in (A). Error bars represent SEM from 3 independent experiments with >30 cilia per condition. Significance was determined by one-way ANOVA; ****P < 0.0001. (C) NIH-3T3 fibroblasts were treated as in (A) with addition of 0.3 mM Cholesterol:MβCD complex or 80 µM Myriocin for 48h; *Gli1* and *Ptch1* RNA levels were quantified and normalized to a GAPDH control. Error bar represents SEM from 3 independent experiments. Significance was determined by student’s t-test. For *Gli1*, ***p = 0.0003 for DMSO vs. CBE; ***p = 0.0002 for CBE vs. CBE+Chol:MβCD; ***p = 0.0006 for CBE vs. CBE+Myriocin. For *Ptch1*, **p = 0.0021 for DMSO vs. CBE; ***p = 0.0003 for CBE vs. CBE+Chol:MβCD; and ****P < 0.0001 for CBE vs. CBE+Myriocin. (D) Quantification of *Gli1* mRNA, normalized to GAPDH, from wild type or GBA1 D409V heterozygous MEF cells stimulated ± 25nM SAG in medium without serum for 24h. Error bars represent SEM from three independent experiments. Significance was determined relative to the control (WT) by Student’s t-test. ***P = 0.0002. (E,F) NIH-3T3 Fibroblasts were treated ±100 µM CBE for 72h and stained for Arl13b (magenta), a primary cilia marker, to quantify ciliation by direct counting. Bar,10µm. (F) Quantification of percent ciliation of NIH-3T3 fibroblasts as in (E). Error bars represent SEM from three independent experiments with >300 cells per condition. Significance was determined by student’s t-test and was not significant.

Losses in ciliary accessible cholesterol are predicted to cause defects in Hedgehog signaling^27,28^. Indeed, incubation of wild type NIH-3T3 cells with the GBA1 inhibitor CBE significantly decreased their ability to respond to Sonic Hedgehog (Shh) ligand, as monitored by changes in the induction of *Gli1* and *Ptch1* mRNAs (Figure 3C). Decreased Hedgehog signaling paralleled loss of ciliary accessible cholesterol (Figure 3B). Similarly, embryonic fibroblasts from heterozygous *Gba1* D409V mice – which exhibit significantly decreased GCase activity in liver and brain^30^– showed a poor response to a Hedgehog agonist compared with wild type fibroblasts, as monitored by induction of *Gli1* transcripts (Figure 3D). CBE addition caused no change in ciliation in NIH-3T3 cells, as monitored using antibodies against Arl13B found in primary cilia (Figure 3E,F). Thus, loss of ciliary accessible cholesterol in GBA1-inhibited and GBA-deficient cells correlates with decreased Hedgehog signaling, without altering cellular ciliation status.

### Cholesterol addition rescues Hedgehog signaling in GBA1-inhibited cells

If the decrease in Hedgehog signaling seen in GBA1-deficient or GBA1-inhibited cells is truly due to the absence of ciliary accessible cholesterol, it should be possible to rescue this phenotype by addition of exogenous cholesterol. To test this, we added cholesterol to cells as cholesterol-methyl beta-cyclodextrin complexes (Chol:MβCD)^31^. An alternative method to increase plasma membrane accessible cholesterol is to inhibit sphingomyelin biosynthesis by addition of myriocin^27^. Addition of Chol:MβCD increased accessible cholesterol (Figure 3A,B) and treatment with either Chol:MβCD or myriocin fully restored or even enhanced Hedgehog signaling (Figure 3C), despite the continued absence of GBA1 activity. These data strongly support the conclusion that GBA1 inhibition interferes with Hedgehog signaling by decreasing plasma membrane and ciliary accessible cholesterol levels.

### GBA1 inhibition activates the cholesterol biosynthetic pathway

Upon NPC1 mutation or inhibition, cholesterol accumulation in lysosomes and depletion from the plasma membrane triggers the cell’s cholesterol-sensing mechanisms to induce transcription of cholesterol biosynthesis genes^20,21^. If *GBA1* mutation causes lysosomal cholesterol accumulation, one might expect a similar, apparent “cholesterol starvation” transcriptional response. Total sequencing of cellular RNA revealed that GBA1 inhibition does indeed trigger a cholesterol-starved condition as shown in a heatmap of genes induced by CBE treatment (Figure 4A) and mapped onto the cholesterol biosynthetic pathway (Figure 4B). The entire cholesterol biosynthesis pathway was activated, including HMG CoA-Synthase 1, the rate limiting HMG CoA-reductase, the LDL receptor and SREBP2, which senses cholesterol levels in the ER. Independent PCR reactions confirmed induction of representative cholesterol biosynthesis pathway genes (Figure 4C). These data confirm cholesterol metabolic dysregulation in GBA1-inhibited cells.

**Figure 4:**
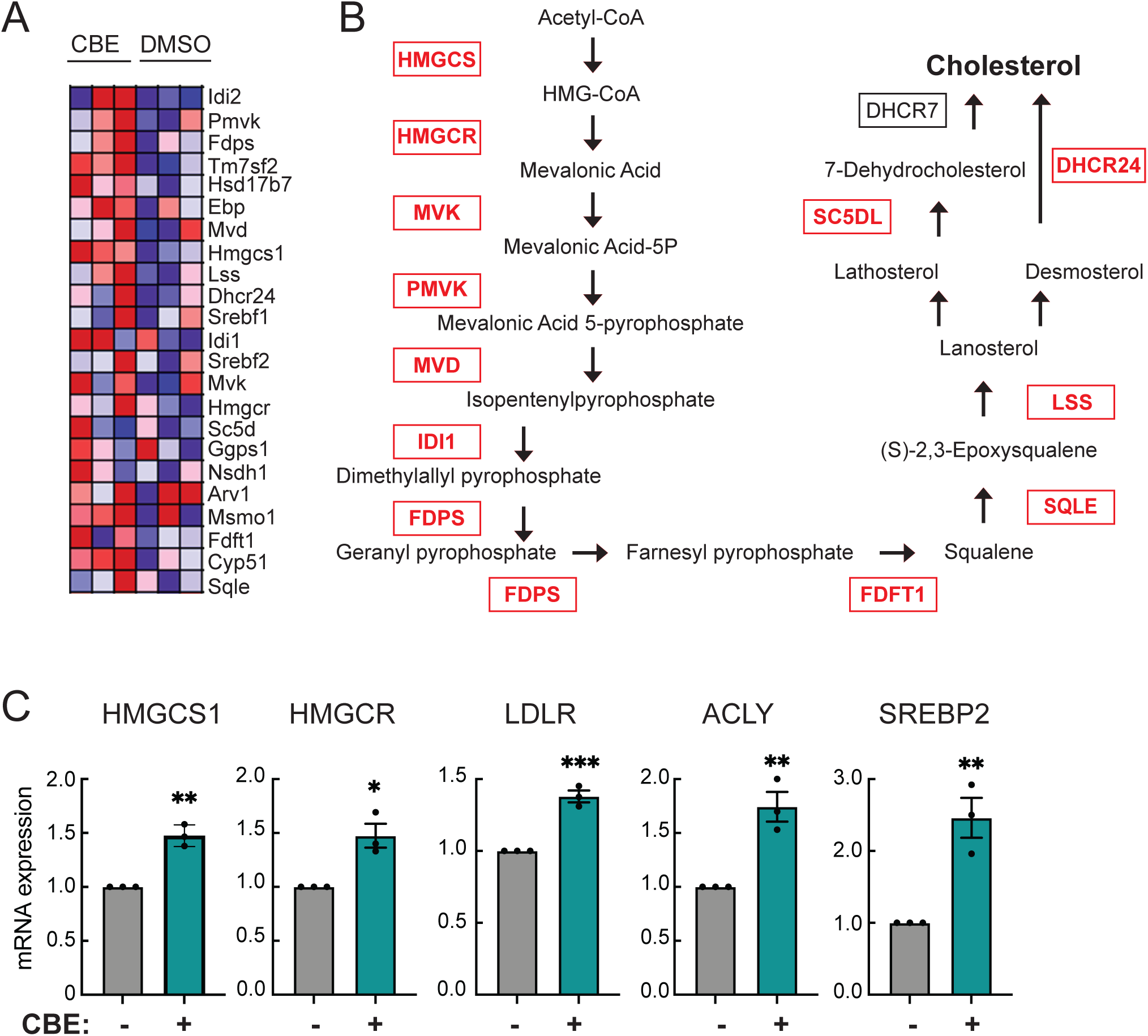
GBA1 inhibition up-regulates cholesterol biosynthesis. (A) Heatmap of enriched genes in DMSO-vs. CBE-treated NIH-3T3 fibroblasts obtained from bulk RNAseq analysis. Each column represents an individual biological sample (n=3 per group). Each row represents individual genes enriched as indicated in the pathway shown in (B). (B) Cholesterol biosynthesis pathway. Enzymes highlighted in red were all upregulated in the RNAseq dataset. (C) qPCR re-validation of key enzymes of cholesterol biosynthesis pathway from NIH-3T3 fibroblasts treated as in (A). Error bars represent SEM from three independent experiments. Significance was determined relative to control, (DMSO, set to 1) by student’s t-test. **p = 0.0012 for HMGCS-1, *p = 0.0129 for HMGCR, ***p = 0.0008 for LDLR, **p = 0.0057 for ACLY, and **p = 0.0062 for SREBP2.

### GBA1 mutations block Hedgehog signaling in the Nigrostriatal circuit

We have shown previously that pathogenic, activating LRRK2 mutations block primary ciliogenesis in certain rare cell types in the dorsal striatum that are important for providing dopamine neuron neuroprotection^11–14^. Specifically, cholinergic and parvalbumin interneurons, but not the more abundant medium spiny neurons, lose primary cilia and therefore fail to respond to Hedgehog signals and fail to induce GDNF and NRTN expression to support dopamine neurons^10–12^. About half of astrocytes in the dorsal striatum also lose cilia and therefore, their ability to respond to Hedgehog signals^14^.

The experiments presented above show that *Gba1* mutations do not alter primary ciliation in cell culture, but nevertheless decrease ciliary accessible cholesterol, and thus interfere with Hedgehog signaling. Therefore, we sought to determine if such a Hedgehog signaling blockade and loss of neuroprotection could also be detected in *Gba1*-mutant mouse brain. For these experiments we analyzed brains from homozygous *Gba1* D409V mice. This mouse model has been shown to exhibit significantly decreased GCase activity in liver and brain^30^.

Cholinergic neurons in the dorsal striatum were identified using anti-choline acetyltransferase antibodies; astrocytes were identified using anti-Aldh1l1 antibodies, and their primary cilia were labeled with anti-adenylate cyclase 3 (AC3) or Arl13B, respectively. Consistent with cells in culture, *GBA1* mutant cholinergic neurons and astrocytes were ciliated to the same extent as those in wild type animals (Figure 5A-D).

**Figure 5:**
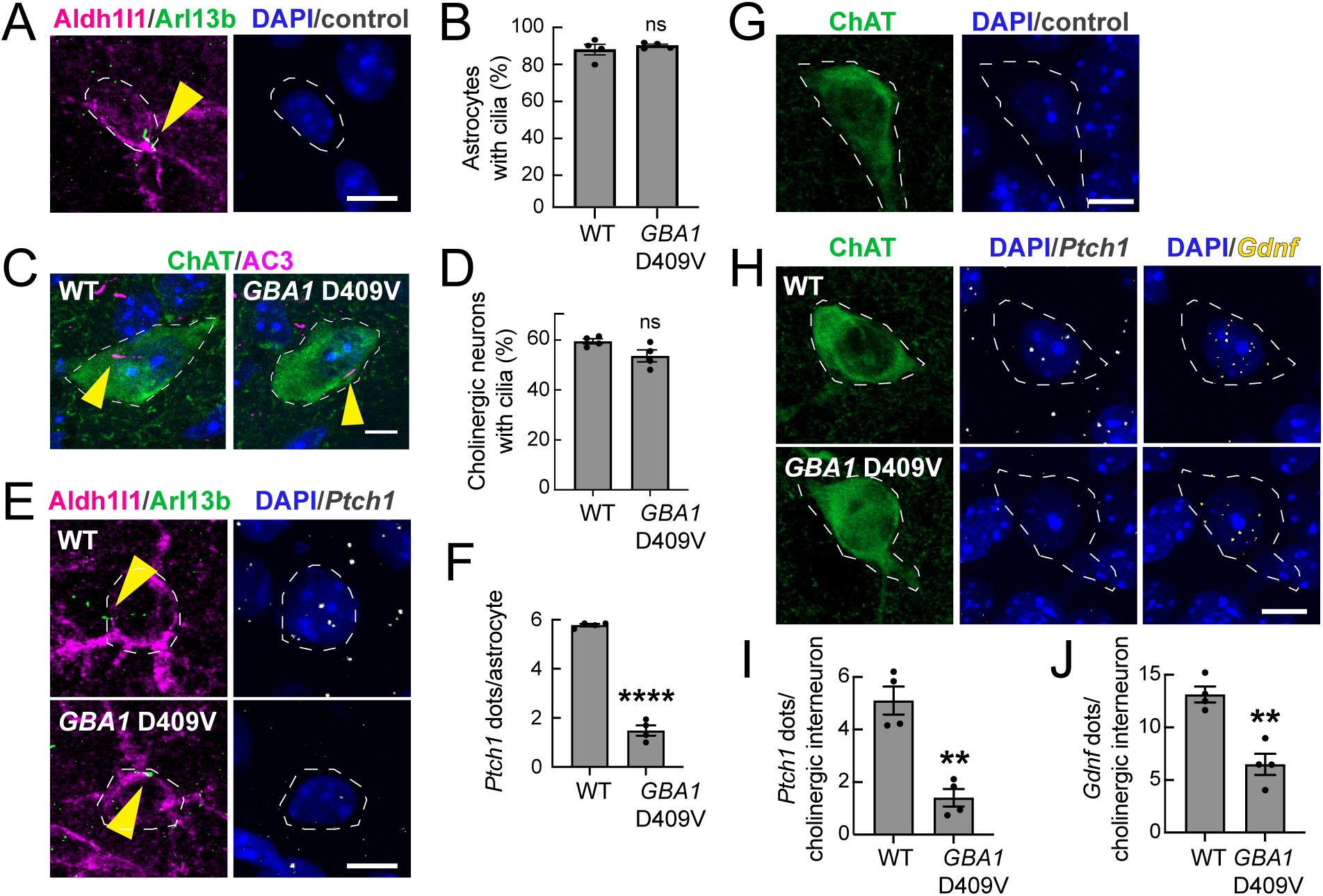
Decreased *Ptch1* and *Gdnf* expression in *GBA1* mutant mouse striatal cholinergic interneurons and astrocytes. (A) Confocal images of dorsal striatal control reactions carried out without the RNAscope in situ hybridization probe in wild type (WT) mouse striatal astrocytes. Astrocytes were identified using anti-ALDH1L1 antibody (magenta); primary cilia were labeled using anti-Arl13b antibody (green); nuclei were labeled using DAPI (blue). (B) Percentage of astrocytes with cilia in 3-month-old WT and *Gba1* D409V homozygous animals (no significant difference). (C) Example images of striatal cholinergic neurons (labeled with anti-choline acetyltransferase/ChAT antibody) and their primary cilia (labeled with anti-adenylate cyclase 3/AC3 antibody) from WT and *Gba1* D409V homozygous animals. (D) Percentage of cholinergic interneurons with cilia. Error bars represent the mean ± SEM from 4 WT and 4 *Gba1* D409V homozygous brains, each containing >25 cells. Significance was determined by unpaired student’s t-test and was not significant. (E) WT and *Gba1* D409V homozygous mouse dorsal striatum were subjected to in situ hybridization using *Ptch1* RNA probe (white dots) in astrocytes. (F) Quantification of average number of *Ptch1* dots per ALDH1L1^+^ astrocyte. Error bars represent the mean ± SEM from 4 WT and 4 D409V HOM brains, each containing >40 cells. Significance was determined by unpaired student’s t-test and ****P < 0.0001. (G) Confocal images of control reactions carried out without the RNAscope probe in WT mouse striatal cholinergic interneurons. (H) Brain sections as in (G) were subjected to in-situ hybridization using *Ptch1* (white dots) and *Gdnf* (yellow dots) RNA probes. Cholinergic interneurons (ChAT^+^) were identified using anti-ChAT antibody (green); nuclei (blue) were stained with DAPI. (I) Quantification of *Ptch1* dots per ChAT^+^ neuron and (J) average number of *Gdnf* dots per ChAT^+^ neuron. Error bars represent SEM from 4 GBA1 WT brains and 4 GBA1 D409V homozygous brains with >40 ChAT^+^ neurons scored per brain. Significance was determined by unpaired student’s t-test. For *Ptch1* dots, **p = 0.0011; for *Gdnf* dots, **p = 0.0019. Bar,10µm.

To measure expression of Hedgehog pathway genes in brain tissue, we used a fluorescence in situ hybridization method, RNAscope^TM^ (Advanced Cell Diagnostics). We monitored Patched1 (*Ptch1*) RNA, a reliable Hedgehog-induced gene that encodes the receptor for Shh, as well as *Gdnf* RNA, to determine the consequences of *GBA1* mutation on dopaminergic neuroprotection.

Comparison of wild type and *GBA1*-D409V brain astrocytes revealed a dramatic decrease in expression of *Ptch1* RNA in the *GBA1* mutant (Figure 5E-F). Control reactions carried out in the absence of the *Ptch1* RNAscope^TM^ probe showed no RNA signal (Figure 5A). These results suggest that, despite possessing cilia, GBA1-mutant astrocytes do not respond to Hedgehog ligand to induce *Ptch1* RNA synthesis. Note that the major decrease in *Ptch1* expression is predicted to also decrease the ability of these cells to further sense and respond to Hedgehog signals.

Similar to our results in astrocytes, we found that cholinergic interneurons also had decreased *Ptch1* expression in the *GBA1* D409V mutant dorsal striatum (Figure 5G-I). Moreover, these neurons also showed an ∼50% decrease in their *Gdnf* RNA content, as measured by RNAscope^TM^ in situ hybridization (Figure 5G,H,J), further supporting the conclusion that Hedgehog signaling is decreased. These data support the hypothesis that dysregulation of accessible cholesterol blocks the ability of these ciliated cells to carry out a normal Hedgehog signaling response that is needed to induce *Gdnf* transcription in *GBA1* mutant striatum.

In the context of PD, Hedgehog signaling is critical for neuroprotection of dopamine neurons: *Lrrk2* mutation leads to loss of *Gdnf* and GDNF-related *Nrtn* expression upon cilia loss by cholinergic and parvalbumin interneurons, respectively^11,12^. *Lrrk2* mutation also interferes with ciliation of Aldh1l1- and GFAP-expressing astrocytes, and we find here that this correlates with loss of Brain-derived neurotrophic factor (BDNF) expression in these astrocytes in both *Lrrk2* mutant astrocytes (Figure 6A-C) and *Gba1* mutant astrocytes (Figure 6D-F). In both genetic backgrounds, the expression of *Bdnf* was highest in ciliated cells and lower in mutant cells, whether or not the mutant cells were ciliated (Figure 6 C,F). These data confirm general Hedgehog signaling defects in both *Lrrk2* and *Gba1* mutant striatum and add an additional neurotrophic factor that may contribute to PD upon its loss.

**Figure 6:**
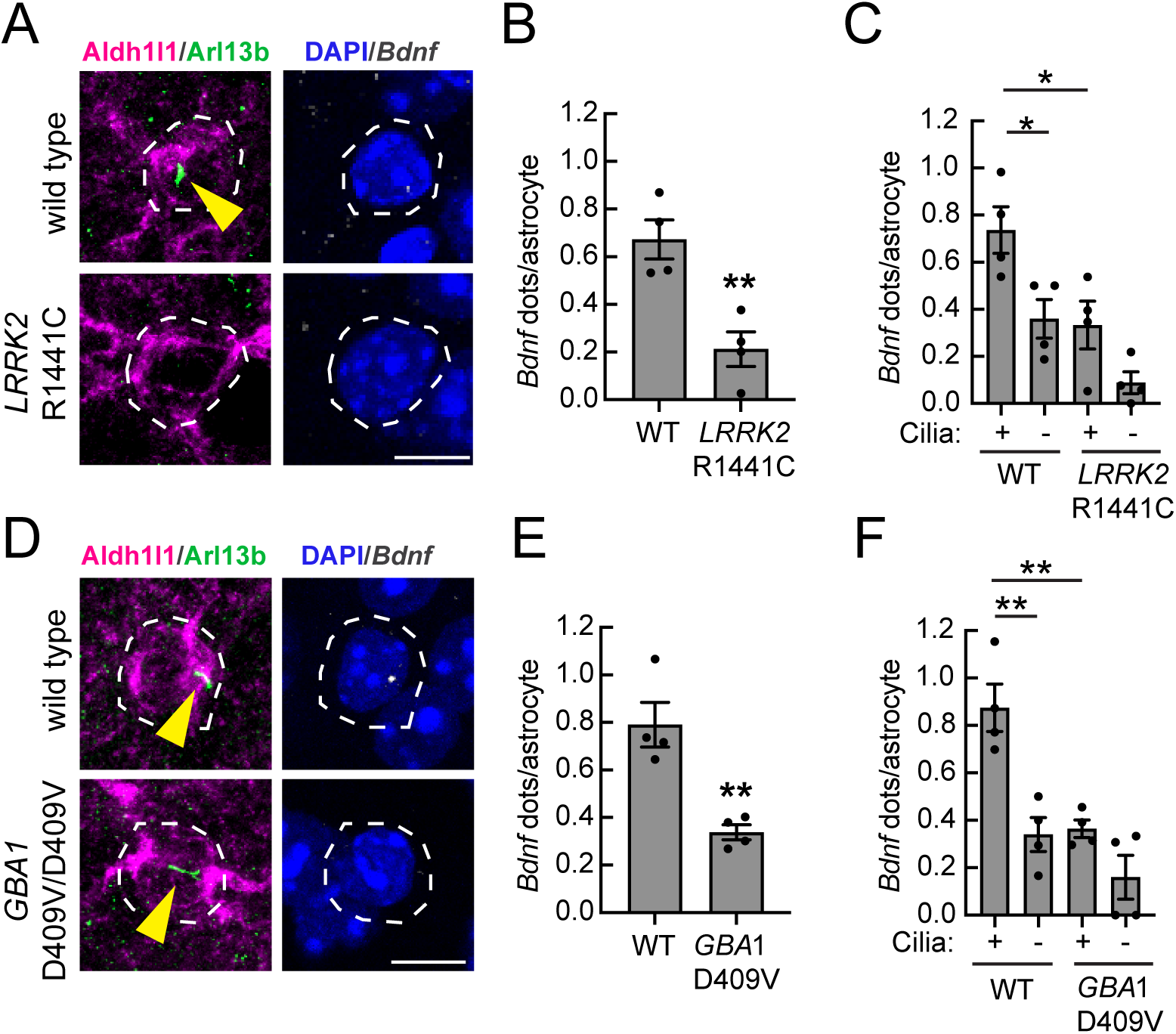
Decreased *Bdnf* expression in *LRRK2* and *GBA1* mutant mouse striatal astrocytes. (A) Confocal images of Aldh1l1^+^ astrocytes from 5-month-old WT or R1441C *LRRK2* mouse dorsal striatum subjected to in-situ hybridization using *Bdnf* RNA probe (white dots). Astrocytes were identified using anti-ALDH1L1 antibody (magenta); primary cilia were labeled using anti-Arl13b antibody (green); nuclei were labeled using DAPI (blue). (B) Quantification of *Bdnf* dots per ALDH1L1^+^ astrocyte (C) or same data segregated as a function of ciliation status. Error bars represent the mean ± SEM from four WT and four R1441C LRRK2 brains, each containing >60 astrocytes. Significance was determined by unpaired student’s t-test or one-way ANOVA. **P = 0.0056 for WT versus R1441C LRRK2; *P = 0.0379 for WT ciliated versus WT unciliated; *P = 0.0258 for WT ciliated versus R1441C LRRK2 ciliated. (D) Similar analysis of 3-month-old WT and *Gba1* D409V/D409V homozygous animals subjected to in-situ hybridization using *Bdnf* RNA probe (white dots). (E) Quantification of *Bdnf* RNA dots per ALDH1L1^+^ astrocyte (F) or same data segregated as a function of ciliation status. Error bars represent the mean ± SEM from 4 WT and 4 *Gba1* D409V homozygous brains, each containing >55 astrocytes. Significance was determined by unpaired student’s t-test or one-way ANOVA. **P = 0.0038 for WT versus *Gba1* D409V homozygous; **P = 0.0022 for WT ciliated versus WT unciliated; **P = 0.0032 for WT ciliated versus *Gba1* D409V homozygous ciliated. Bar, 10µm.

## DISCUSSION

We have shown here that *Gba1* mutations or GBA1 chemical inhibition interferes with Hedgehog signaling in cultured cells. Moreover, GBA1 inhibition decreases the availability of accessible cholesterol in primary cilia that is a requirement for such signaling. Indeed, GBA1 inhibition leads to accumulation of accessible cholesterol in lysosomes and triggers upregulation of the cholesterol biosynthetic pathway, as determined by total cell RNA sequencing. Finally, in critical cell types of the brains of mice with *Gba1* mutations, we detect major decreases in Hedgehog signaling, leading to the specific loss of neuroprotective *Gdnf* RNA production by cholinergic neurons and *Bdnf* RNA expression in astrocytes of the dorsal striatum. These changes are reminiscent of those seen in *Lrrk2* mutant mice^11–13,15^ and provide a plausible explanation for how *Gba1* mutations may increase PD risk in a manner similar to that seen upon LRRK2 mutation^11^: vulnerable neurons fail to receive adequate neuroprotection from cholinergic interneurons and present with a disease of aging.

In humans with *LRRK2* mutations, about 50% of interneuron cilia are lost^11^ and in mice, cilia loss is accompanied by loss of *Ptch1* transcripts. In patients with *GBA1* mutations, our data suggest that cilia will be signaling-defective, but subtly so, such that patients show no developmental defects and present with PD vulnerability late in life. In a patient with both *LRRK2* and *GBA1* mutations, specific cell types will have fewer cilia and the remaining cells may upregulate Shh signaling components to compensate for losses of cilia on other neurons. Indeed, we have detected anomalous *Gli1* transcription in cells that retain their cilia, despite *LRRK2* mutation^13,14^.

A hallmark of PD is loss of dopamine neurons in the Substantia nigra. We show here that multiple genetic causes of PD–i.e. *LRRK2* mutation^11–13^, *GBA1* mutation (this study), as well as loss of PINK1 function^32^ all cause decreases in the production of RNAs encoding critical neuroprotective factors, GDNF, NRTN, and now, shown here, BDNF, due to impaired Shh signaling. In the case of LRRK2 hyperactivation or *PINK1* knockout, loss and/or shortening of cilia in specific cell types blocks Hedgehog signaling. Here, we found that *GBA1* mutation does not alter overall ciliation but nevertheless, interferes with Hedgehog signaling due to altered ciliary lipid composition. We have also shown recently that primary cilia are lost from cholinergic neurons, but not medium spiny neurons of the lenticular nucleus (striatal equivalent) in postmortem tissue from patients with idiopathic PD^11^. Why these cilia are lost in idiopathic disease will be an important question for future study, but a certain consequence will be loss of Hedgehog signaling.

Idiopathic PD is considered a type of Lewy body disease, where synuclein aggregates accumulate in and around lysosomes and lead to cell death. GBA1 activity is lower in brain tissue of sporadic PD patients with synuclein pathology as compared to controls^33,34^. Decreased GBA1 activity can decrease synuclein aggregate clearance through impairment of lysosome function and accumulation of GBA1 substrates that promote synuclein aggregation ^33,35–38^. It is very possible that such loss of GBA1 activity influences the Hedgehog signaling capacity of synuclein aggregate-containing neurons. Note that autophagy is linked to ciliogenesis^39,40^, thus lysosome dysfunction caused by Lewy Body formation is likely to also impact Shh signaling.

We have shown here that astrocytic cilia loss in *Lrrk2* mutant mice decreases their production of *Bdnf* RNA, and cilia loss by cholinergic and Parvalbumin neurons decreases striatal *Gdnf* and *Nrtn* RNAs^11,12^ that are critical for the survival, differentiation, and function of dopaminergic neurons. *Gba1* or *Lrrk2* mutation decreases *Bdnf* production by striatal astrocytes, as would be expected from a general block in Hedgehog signaling.

Recent single nucleus RNA sequencing experiments have identified a subpopulation of dopamine neurons that are specifically vulnerable in PD^41,42^. These neurons represent a subset of Aldh1a1/Sox6^+^ neurons that also express Anxa1; they project into the dorsolateral striatum where Hedgehog signaling is most robust^41,42^. We predict that these neurons will be the most sensitive to decreases in Hedgehog signaling and it will be important to use conditional expression of *Lrrk2* or *Gba1* mutations to explore the relative contribution of striatal Hedgehog signaling to Anxa1/Aldh1a1/Sox6+ neuronal vulnerability.

Finally, recent work has shown that neuronal primary cilia are likely to be more important for circuit regulation than previously believed^43,44^. In the adult brain cortex, interneuron cilia may contact as many as 40 different neurons via projections that seem to congregate over ciliary structures. These projections often form synapse-like connections and in some cases, apparent gap junctions are also detected^44^. Thus, decreased ciliary signaling in the striatum in *LRRK2* or *GBA1* mutant carriers and idiopathic PD will very likely have broader consequences for overall neuronal signaling in the striatum.

## Resource availability

Lead Contact: Suzanne Pfeffer

All reagents used in this study are available from commercial sources or repositories without restriction and RRIDs for all reagents are provided in the Key Resources Table. All primary data are available at zenodo.org/records/12753029.

## Acknowledgments

This study was funded by the joint efforts of The Michael J. Fox Foundation for Parkinson’s Research (MJFF) and Aligning Science Across Parkinson’s (ASAP) initiative. MJFF administers the grant (ASAP-000463) on behalf of ASAP and itself (to SRP). Work on NPC1 was supported by a grant from Notre Dame University and the Ara Parseghian Medical Research Foundation. We thank Rajat Rohatgi lab and Arun Radhakrishnan for gifts of Sonic Hedgehog protein and mNeon-ALOD4 plasmid, respectively, Mirella Bucci for comments on the manuscript and Maia Kinnebrew and Kathy Le for helpful discussions.

For the purpose of open access, the authors have applied for a CC-BY public copyright license to the Author Accepted Manuscript version arising from this submission.

## Author contributions

SVN and EJ carried out all experiments; AA purified invaluable mNeon-ALOD4 protein and JN helped quantify ciliary accessible cholesterol. SRP directed the project, acquired funding and wrote the manuscript.

## Declaration of generative AI and AI-assisted technologies

No generative AI and AI-assisted technologies were employed in the writing process.

## Key Resource Table

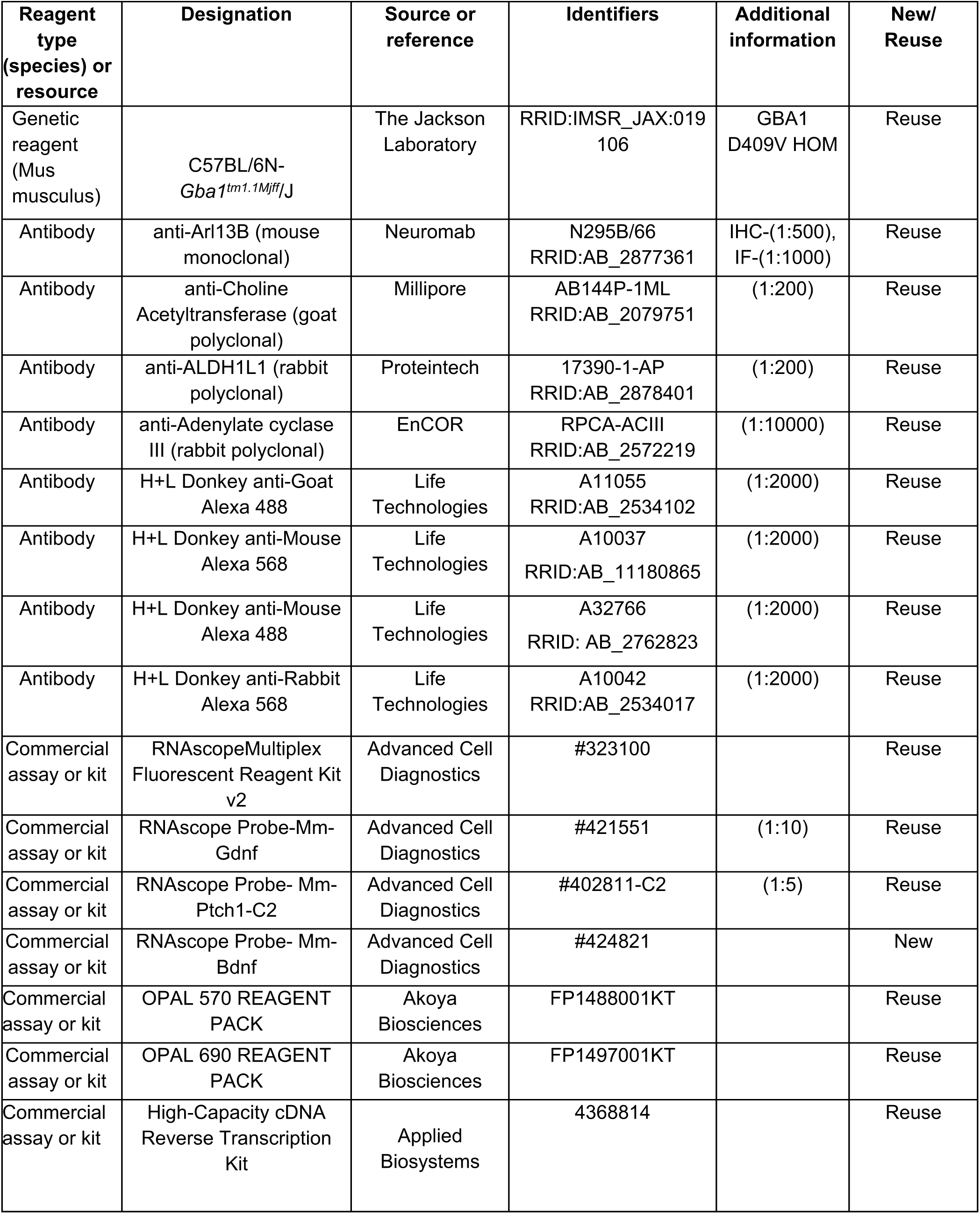
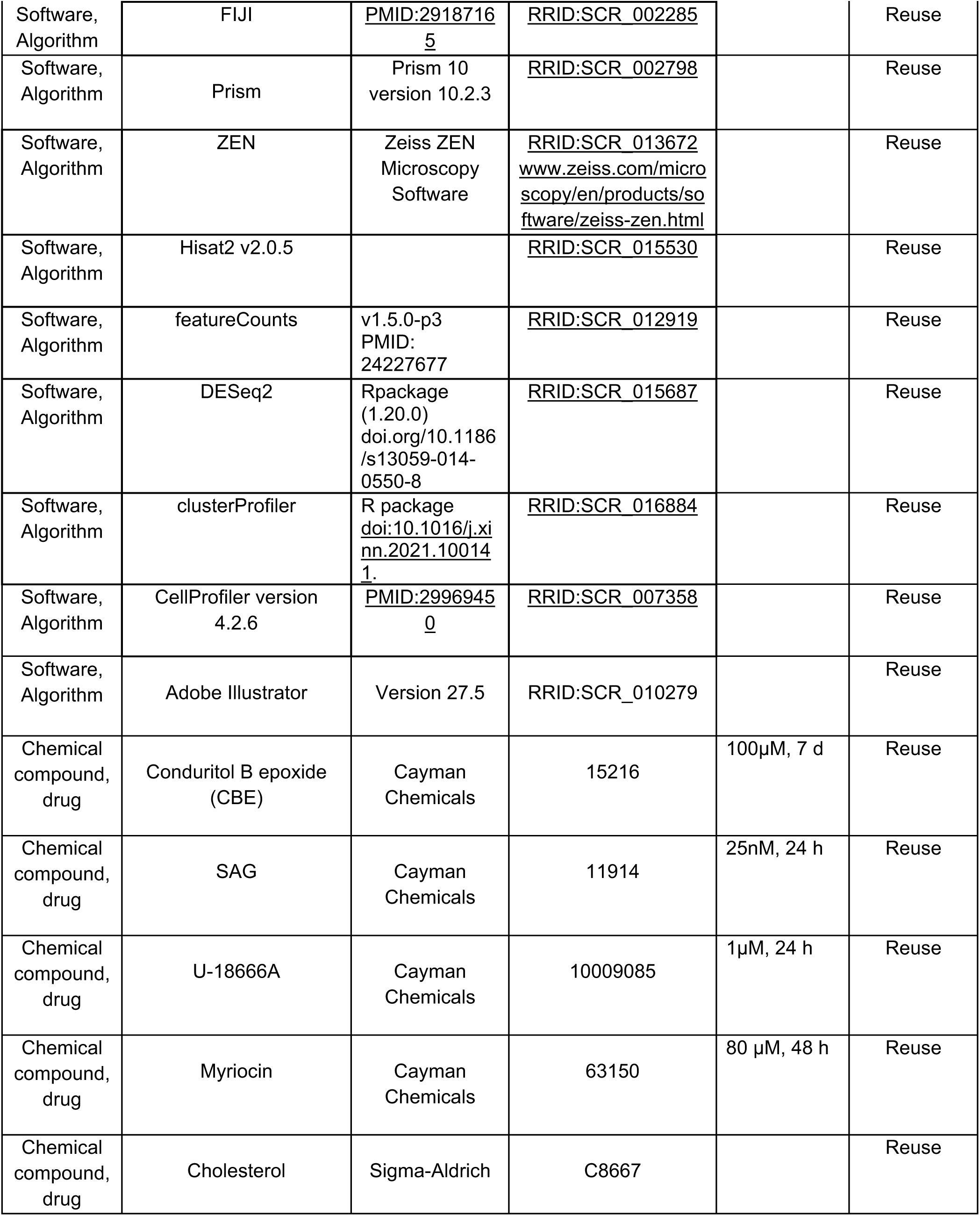
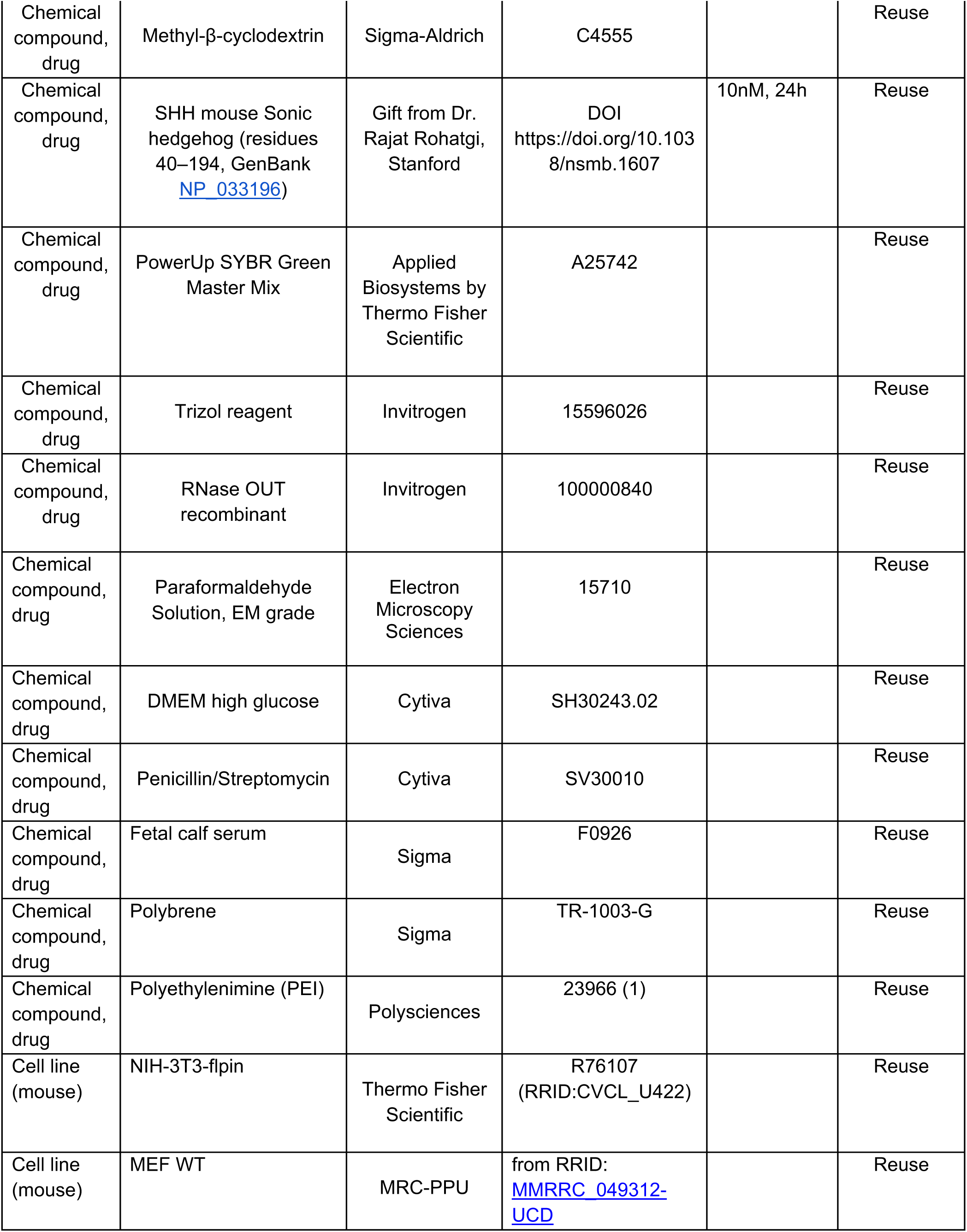
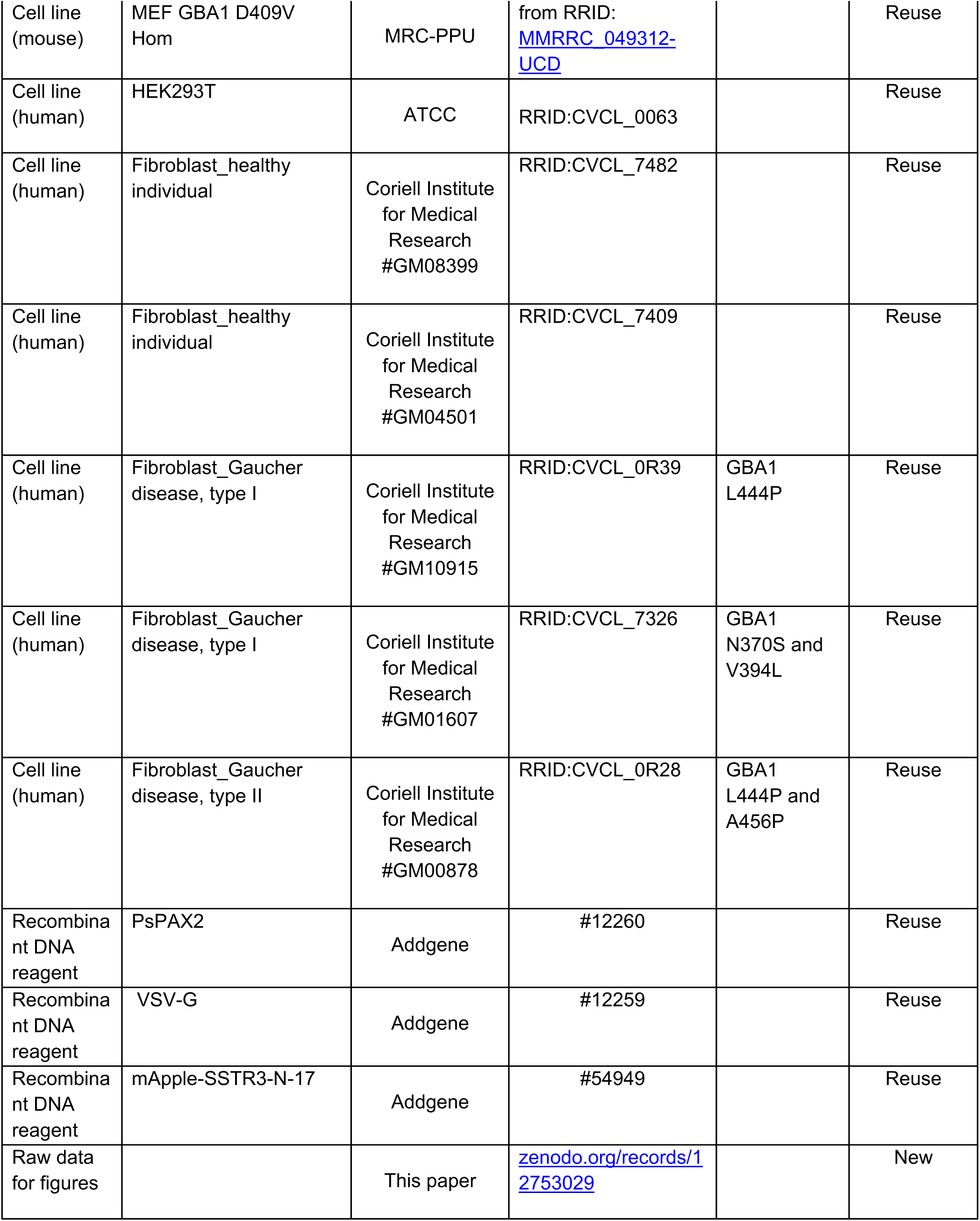
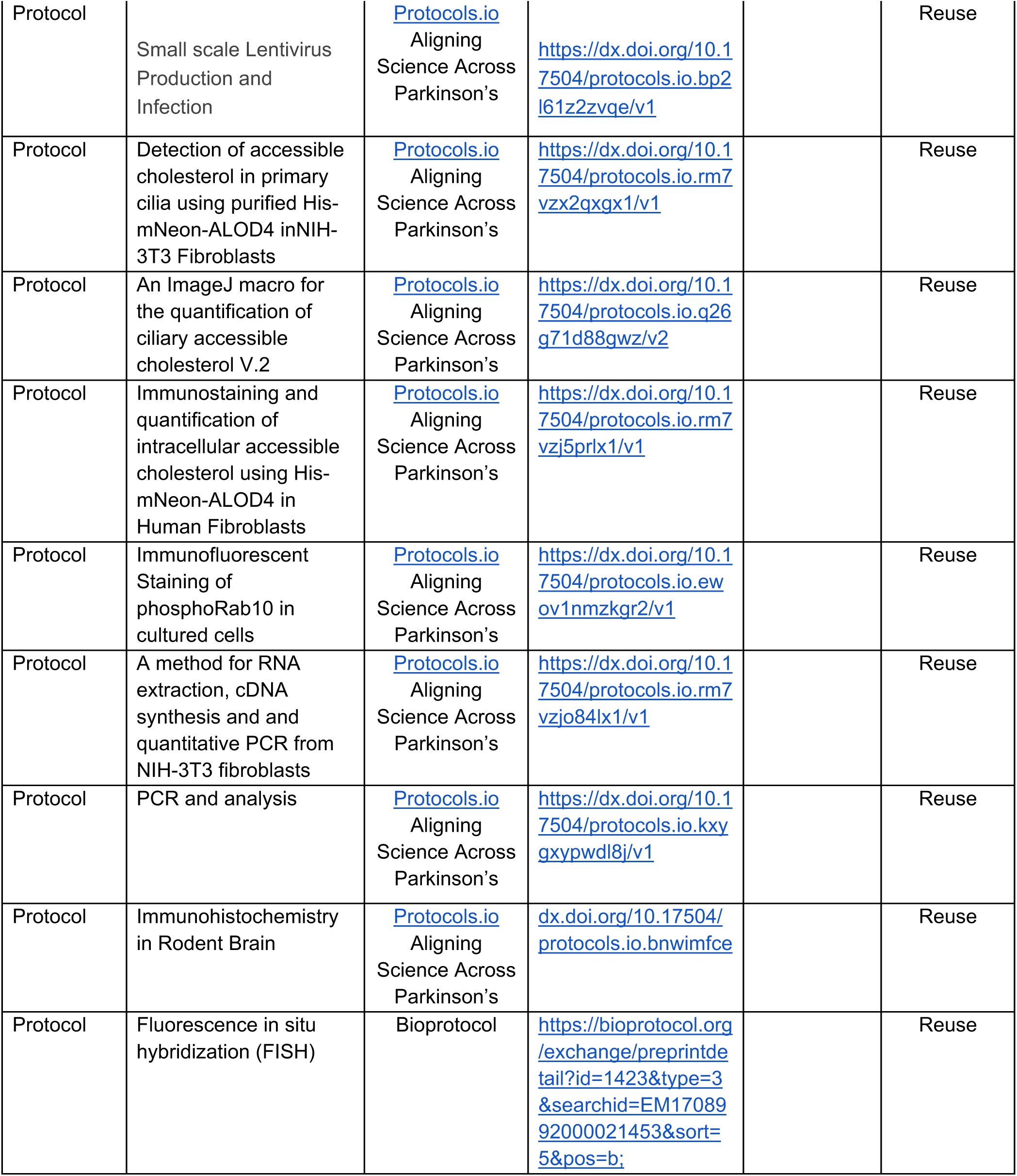

## STAR Methods

### Cell Culture

NIH-3T3, MEF, and HEK293T cells were cultured in high-glucose DMEM containing 10% (v/v) fetal calf serum, 2 mM L-glutamine, 100 U/mL penicillin, and 100 µg/mL streptomycin. All cells were grown at 37°C, 5% CO_2_ in a humidified atmosphere and regularly tested for *Mycoplasma* contamination. To generate cells stably expressing SSTR3-mApple, HEK293T cells were transfected with CSII-SSTR3-mApple (1μg), PsPAX2 (1μg) and VSV-G (0.5μg) using 1 mg/mL polyethylene imine (PEI) at ratio 5:1 (PEI:DNA). A detailed protocol can be found here: (https://dx.doi.org/10.17504/protocols.io.bp2l61z2zvqe/v1). After 72h, the supernatant containing viral particles was collected and concentrated by ultracentrifugation at 90000 x g, 90 min at 4°C. 0.2 mL of concentrated virus was used to infect NIH-3T3 cells in one well of a 6-well plate along with 5µg/mL Polybrene for 48h. These cells were grown in complete media for a week and selected for SSTR3-mApple expression by flow cytometry.

### Protein Purification

His-mNeon-FLAG-ALOD4 protein was purified after expression in E. coli BL21 (DE3 pLys), as described (https://dx.doi.org/10.17504/protocols.io.rm7vzx2qxgx1/v1). His-mNeon-FLAG-ALOD4 plasmid was a gift from Arun Radhakrishnan. Bacterial cultures were grown at 37°C in Luria Broth and induced with 0.3 mM isopropyl-1-thio-β-d-galactopyranoside overnight at 18°C to an OD 600 nm reached 0.5-0.6. Cell pellets were resuspended in ice-cold lysis buffer (50 mM HEPES, pH 8.0, 10% glycerol, 500 mM NaCl, 10 mM imidazole, 5 mM MgCl_2_, 0.2 mM TCEP, 20 μM GTP, and EDTA-free protease inhibitor cocktail (Roche #69040300). The bacteria were lysed using an Emulsiflex-C5 apparatus (Avestin) at 10,000 psi and centrifuged at 40,000 rpm for 45 minutes at 4°C using a Beckman Ti45 rotor. The cleared lysate was filtered through a 0.2 µm filter (Nalgene) and passed over a HiTrap TALON crude 1 mL column (Cytiva #28953766). The column was washed with lysis buffer until the absorbance returned to baseline; protein was eluted with a gradient of 100 to 500 mM imidazole in lysis buffer. Peak fractions were identified by SDS-PAGE on 4-20% Precast Protein Gels, and protein was buffer exchanged using a PD-10 desalting column (Cytiva #17085101) with 50 mM HEPES, pH 8, 10% glycerol, 150 mM NaCl, 5 mM MgCl_2_, and 0.1 mM TCEP. Protein was used fresh or stored at 4°C for up to two weeks after purification as ALOD4 loses its activity if frozen.

### mNeon-ALOD4 staining

Detection of ciliary accessible cholesterol was carried out as described: (https://dx.doi.org/10.17504/protocols.io.rm7vzx2qxgx1/v1). Briefly, NIH-3T3 cells expressing SSTR3-mApple were treated with 100 μM conduritol beta epoxide (CBE) or DMSO (vehicle control) for 7d. On the 6^th^ day of treatment, cells were starved in media without serum for 24h in the presence of DMSO or CBE to induce ciliogenesis. Cells were fixed with 4% (v/v) paraformaldehyde for 10 min at room temperature (RT). Cells were washed 3X with 1X PBS and transferred to an incubation chamber on ice. Coverslips were incubated with 4μM mNeon-ALOD4 in 1% BSA for 1h on ice. Cells were washed twice in 1X PBS and fixed with 2% (v/v) paraformaldehyde for 5 min at RT. Coverslips were washed twice in 1X PBS and mounted onto clean glass slides using 4μL Mowiol. Slides were stored at RT in dark until imaging. Images were obtained using a Zeiss LSM 900 confocal microscope with a 63 X 1.4 oil immersion objective. Images were processed as described in this protocol https://dx.doi.org/10.17504/protocols.io.q26g71d88gwz/v2 for the quantification of ciliary accessible cholesterol.

Immunostaining and quantification of intracellular accessible cholesterol has been described in detail: https://dx.doi.org/10.17504/protocols.io.rm7vzj5prlx1/v1. Briefly, human fibroblasts were fixed with 4% (v/v) paraformaldehyde for 15 min, permeabilized with 0.1% saponin for 5 min, and blocked with 1% BSA in 1X PBS for 30 min at RT. Coverslips were incubated with 4μM mNeon-ALOD4 in 1% BSA in 1X PBS for 1h at RT. Cells were washed thrice with 1X PBS and mounted onto clean glass slides using 4μL Mowiol. Slides were stored at RT in dark until imaging.

### Immunofluorescence microscopy

NIH-3T3 cells were treated with 100 μM CBE or DMSO for 72h. After 48h, cells were transferred to media without serum in the presence of DMSO or CBE to induce ciliogenesis for 24h. After 72h, cells were fixed with 4% (v/v) paraformaldehyde for 10 min, permeabilized with 0.1% Triton X-100 for 10 min, and blocked with 1% BSA for 30 min, all at room temperature as described: (https://dx.doi.org/10.17504/protocols.io.ewov1nmzkgr2/v1). Cells were subsequently stained with mouse anti-Arl13b antibody (Neuromab, N295B/66, 1:1000) for 2h at RT. Cells were washed 3X with PBS and incubated with H+L Donkey anti-Mouse Alexa 568 secondary antibody (Life Technologies, A10037, 1:2000) for 1h at room temperature. Nuclei were stained with 0.1 µg/mL DAPI (Sigma). Images were obtained using a Zeiss LSM 900 confocal microscope with a 63 × 1.4 oil immersion objective. Images were converted to maximum intensity projections using Fiji (https://fiji.sc/) and cilia were counted manually.

### Analysis of Hedgehog signaling and qPCR

NIH-3T3 cells were treated with 10nM recombinant Sonic Hedgehog^45^ for 24 hours in serum free medium to trigger ciliogenesis. In some mouse embryonic fibroblast experiments, SAG Hedgehog agonist was added at 25nM for 24hr in serum free medium. Total RNA extraction was performed as detailed in this protocol (https://dx.doi.org/10.17504/protocols.io.rm7vzjo84lx1/v1). Briefly, media was aspirated from each well of a 6-well plate and 0.5 mL of Trizol reagent (Invitrogen) was added for 5 min at RT. Cells were collected into a fresh 1.5 mL tube and 100 μL chloroform was added, vortexed for 10-15 sec, and centrifuged at 14,000g for 15 min at 4^0^C. The aqueous layer was collected into a fresh tube and mixed with an equal volume of isopropyl alcohol. These samples were centrifuged at 13,000 x g for 10 min at RT. The supernatant was removed, 0.5 mL of 80% ethanol was added to each tube, and centrifuged at 13,000 x g for 10 min at RT. This step was repeated. The supernatant from the final run was removed and the pellet was allowed to air-dry. Air dried RNA pellets were resuspended in 50 μL sterile, RNAase free water. cDNA synthesis was performed using 500ng RNA using a high-capacity cDNA reverse transcription kit (Applied Biosystems) following the manufacturer’s instructions. The cDNA was diluted 20 fold before qPCR and GAPDH was used as an internal control. For qPCR reactions carried out in quadruplicate, 3 µl cDNA was used as template in the PowerUp SYBR green 2X master mix (Applied Biosystems) with 0.2 μM each forward and reverse primers in a qPCR machine (ViiA7 Applied Biosystems). Gene expression was analyzed by the ΔΔCt method. A detailed protocol can be found here https://dx.doi.org/10.17504/protocols.io.kxygxypwdl8j/v1. The primers used for the qPCR were as follows – Gli1-Fw-5’ CCAAGCCAACTTTATGTCAGGG 3’, Rv-5’ AGCCCGCTTCTTTGTTAATTTGA 3’; GAPDH-Fw-5’ AGTGGCAAAGTGGAGATT 3’, Rv-5’ GTGGAGTCATACTGGAACA 3’, HMGCS1-Fw-5’ AGTGTTCTCTTACGGTTCTG 3’, Rv-5’ AAGCCTTGATTTAAGGTCAC 3’; HMGCR-Fw-5’ GATAGCTGATCCTTCTCCTC 3’, Rv-5’ ATGCTGATCATCTTGGAGAG 3’; LDLR-Fw-5’ CATCTTCTTCCCTATTGCAC 3’, Rv-5’ ATGCTGTTGATGTTCTTCAG 3’, ACLY-Fw-5’ AGGAAGTGCCACCTCCAACAGT 3’, Rv-5’ CGCTCATCACAGATGCTGGTCA 3’; SREBP2-Fw-5’ TTCTCTCCCTATTCCATTGAC 3’, Rv-5’ AAGGTGAGGACACATAAGAG 3’; PTCH1-Fw-5’ GAAGCCACAGAAAACCCTGTC 3’, Rv-5’ GCCGCAAGCCTTCTCTAGG 3’.

### Bulk RNA seq analysis

Total RNA from NIH-3T3 fibroblasts cells was extracted as described above (https://dx.doi.org/10.17504/protocols.io.rm7vzjo84lx1/v1). Quantification of isolated RNA was done on a Nanodrop (NanoDrop One, Thermoscientific) and the samples were sent to Novogene, USA for bulk-RNA sequencing and analysis. RNA sample QC, directional mRNA library preparation by PolyA enrichment were performed and these libraries were then sequenced on NovaSeq PE150 platform at Novogene. Hisat2 v2.0.5 was used to align clean reads to mm39 mouse reference genome, and raw count of each sample was evaluated by featureCounts^46^ v1.5.0-p3. Fragments per kilobase of transcript sequence per millions (FPKM) of each gene was calculated based on the length of the gene and reads count mapped to this gene. Differential expression analysis was performed using the DESeq2Rpackage^47^ (1.20.0). The P-values were adjusted using the Benjamini and Hochberg’s approach for controlling the false discovery rate. Genes with an adjusted P-value <=0.05 found by DESeq2 were assigned as differentially expressed. For Gene Ontology (GO)^48,49^ enrichment analysis of differentially expressed genes, the clusterProfiler R package^50^ was used. GO terms with corrected P-value less than 0.05 were considered significantly enriched by differential expressed genes.

### Mouse brain processing

Homozygous GBA1 D409V and age-matched wild type control brains were processed as described in dx.doi.org/10.17504/protocols.io.bnwimfce. After fixing brains by transcardial perfusion using 4% paraformaldehyde (PFA) in PBS, whole brains were extracted, post-fixed in 4% PFA for 24h and then immersed in 30% (w/v) sucrose in PBS until the tissue settled to the bottom of the tube (∼48h). Prior to cryosectioning, brains were embedded in cubed-shaped plastic blocks with OCT (BioTek, USA) and stored at −80 °C. OCT blocks were allowed to reach −20 °C for ease of sectioning. The brains were oriented to cut coronal sections on a cryotome (Leica CM3050S, Germany) at 16–25 µm thickness and positioned onto SuperFrost plus tissue slides (Thermo Fisher, USA).

### Immunohistochemical staining

The mouse brain striatum was subjected to immunostaining following a previously established protocol (dx.doi.org/10.17504/protocols.io.bnwimfce). Frozen slides were thawed at room temperature for 15 minutes and then gently washed twice with PBS for 5 minutes each. Antigen retrieval was achieved by incubating the slides in 10 mM sodium citrate buffer pH 6.0, preheated to 95°C, for 15 minutes. Sections were permeabilized with 0.1% Triton X-100 in PBS at room temperature for 15 minutes, followed by blocking with 2% FBS and 1% BSA in PBS for 2 hours at room temperature. Primary antibodies were applied overnight at 4°C, and the next day, sections were exposed to secondary antibodies at room temperature for 2 hours. Secondary antibodies used were donkey highly cross-absorbed H + L antibodies conjugated to Alexa 488, Alexa 568, diluted at 1:2000. Nuclei were counterstained with 0.1 μg/ml DAPI (Sigma). Finally, stained tissues were mounted with Fluoromount G and covered with a glass coverslip. All antibody dilutions for tissue staining contained 1% DMSO to facilitate antibody penetration.

### Fluorescence in situ hybridization (FISH)

RNAscope fluorescence in situ hybridization was carried out as described (https://bioprotocol.org/exchange/preprintdetail?id=1423&type=3&searchid=EM1708992000021 453&sort= 5&pos=b;^14^. The RNAscope Multiplex Fluorescent Detection Kit v2 (Advanced Cell Diagnostics) was utilized following the manufacturer’s instructions, employing Mm-Ptch1-C2 (#402811-C2) and Mm-Gdnf (#421951). The Mm-Ptch1-C2, Mm-GDNF probes were diluted 1:5 and 1:10 respectively in dilution buffer consisting of 6X saline-sodium citrate buffer (SSC), 0.2% lithium dodecylsulfate, and 20% Calbiochem OmniPur Formamide. Fluorescent visualization of hybridized probes was achieved using Opal 690 or Opal 570 (Akoya Biosciences). Subsequently, brain slices were subjected to blocking with 1% BSA and 2% FBS in TBS (Tris buffered saline) with 0.1% Triton X-100 for 30 minutes. They were then exposed to primary antibodies overnight at 4°C in TBS supplemented with 1% BSA and 1% DMSO. Secondary antibody treatment followed, diluted in TBS with 1% BSA and 1% DMSO containing 0.1 μg/ml DAPI (Sigma) for 2 hours at room temperature. Finally, sections were mounted with Fluoromount G and covered with glass coverslips.

### Microscope image acquisition

All images were obtained using a Zeiss LSM 900 confocal microscope (Axio Observer Z1/7) coupled with an Axiocam 705 camera and immersion objective (Plan-Apochromat 63x/1.4 Oil DIC M27). The images were acquired using ZEN 3.4 (blue edition) software, and visualizations and analyses were performed using Fiji^51^ and CellProfiler^52^.

In addition to above-mentioned methods, all other statistical analysis was carried out using GraphPad Prism version 10.2.3 for Macintosh, GraphPad Software, Boston, Massachusetts USA, www.graphpad.com.

